# Mapping Histone Modifications in Low Cell Number and Single Cells Using Antibody-guided Chromatin Tagmentation (ACT-seq)

**DOI:** 10.1101/571208

**Authors:** Benjamin Carter, Wai Lim Ku, Qingsong Tang, Jee Youn Kang, Keji Zhao

**Author notes:** These authors contributed equally to this work (Benjamin Carter, Wai Lim Ku).

## Abstract

Modern next-generation sequencing-based methods have empowered researchers to assay the epigenetic states of individual cells. Existing techniques for profiling epigenetic marks in single cells often require the use and optimization of time-intensive procedures such as drop fluidics, chromatin fragmentation, and end repair. Here we describe ACT-seq, a novel and streamlined method for mapping genome-wide distributions of histone tail modifications, histone variants, and chromatin-binding proteins in a small number of or single cells. ACT-seq utilizes a fusion of Tn5 transposase to Protein A that is targeted to chromatin by a specific antibody, allowing chromatin fragmentation and sequence tag insertion specifically at genomic sites presenting the relevant antigen. The Tn5 transposase enables the use of an index multiplexing strategy (iACT-seq), which enables construction of thousands of single-cell libraries in one day by a single researcher without the need for drop-based fluidics or visual sorting. We conclude that ACT-seq present an attractive alternative to existing techniques for mapping epigenetic marks in single cells.

---

Techniques for mapping epigenetic states in individual cells have enhanced our understanding of differentiation and cell-to-cell variation. Multiple single-cell approaches have been developed to map transcriptomes and chromatin organization, and these methods are proving invaluable in fields such as cancer research.^1,2^ Comparatively few methods exist for single-cell profiling of histone tail modifications,^1,3^ which play important roles in epigenetic control of gene expression and development.^4^ Here we describe antibody-guided chromatin tagmentation sequencing (ACT-seq), a technique to assay distributions of epigenetic marks in a small number of cells or thousands of single cells simultaneously.

ACT-seq utilizes Tn5 transposase, which is commonly used to map chromatin accessibility and structure.^5–7^ We fused the N terminus of Tn5 transposase to Protein A (PA) to form a novel fusion protein hereafter referred to as PA-Tnp (**Supplementary Fig. 1**). The PA domain of the fusion protein is first bound to an antibody that is selected to target an epigenetic mark or chromatin-bound protein of interest. The complex is then incubated with permeabilized cells and is guided to chromatin by the associated antibody. After washing away unbound complex, the transposition reaction is initiated by addition of an MgCl2-containing buffer, which results in insertion of sequence tags at sites of bound PA-Tnp. The reaction is terminated by incubation with EDTA and proteinase. The labeled fragments are directly amplified using PCR and sequenced using Illumina HiSeq technology. ACT-seq eliminates procedures including prior chromatin fragmentation, immunoprecipitation, end repair, and adaptor ligation and the entire library preparation takes only 5 to 6 hours, which is much shorter compared to other commonly used methods for mapping histone modifications.

To evaluate the efficiency of ACT-seq, we mapped the distributions of a variety of epigenetic features in HEK293T cells: the histone tail modifications H3K4me1, H3K4me2, H3K4me3, and H2K27ac; the histone variant H2A.Z; and the chromatin-binding protein Brd4 (**Fig. 1a**). Visual inspection using a genome browser revealed highly similar distributions of enrichment in the bulk-cell ACT-seq data relative to published ChIP-seq data sets (**Fig. 1a**).^8,9^ By comparison, little to no enrichment was apparent in the ACT-seq mock IgG sample, indicating that the observed signals were antibody-specific. Analysis of statistically significant peaks of enrichment also revealed strong correlations between the data sets obtained using ACT-seq and ChIP-seq (**Supplementary Fig. 2**). Further, we detected strong average enrichment of H3K27ac, H2A.Z, and Brd4 at transcription start sites (TSS) and enhancer regions in our ACT-seq data (**Fig. 1b,c**) in agreement with published studies on these factors.^10–12^ Taken together, these analyses confirm that ACT-seq and ChIP-seq provide comparable information on enrichment of epigenetic marks in bulk-cell samples.

**Figure 1:**
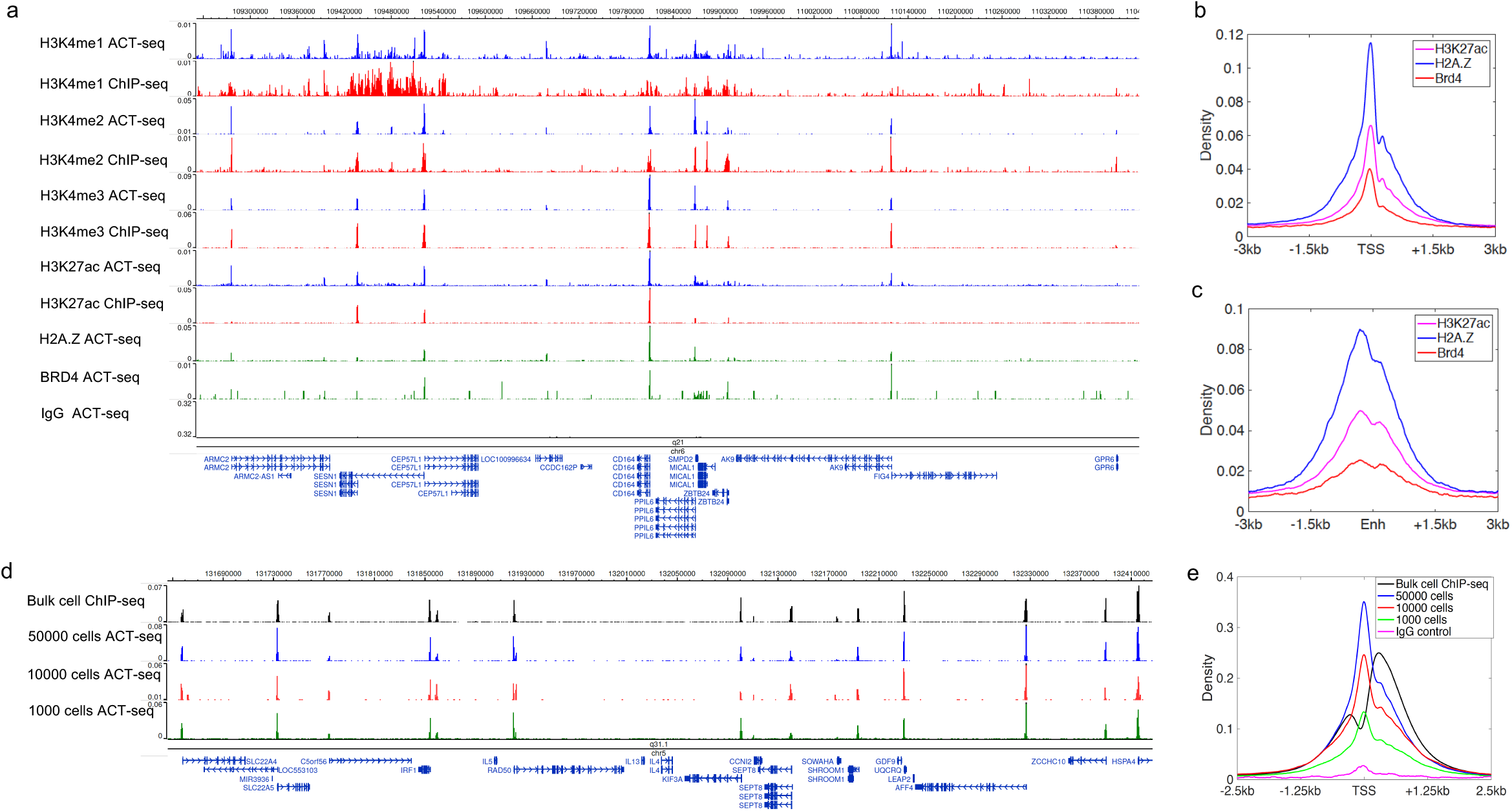
ACT-seq robustly maps epigenetic marks in bulk cell samples. (**a**) Genome browser image depicting enrichment of the indicated epigenetic factors in HEK293T cells at a representative genomic region. Data were obtained using ACT-seq (blue, green) or ChIP-seq (red). The ChIP-seq samples were obtained from published ENCODE data sets. The mock IgG sample is included as a comparative control for enrichment. (**b**) Metagene profile of average H3K27ac, H2A.Z, and Brd4 enrichment at the transcription start site (TSS) region of annotated genes from the hg19 genome. (**c**) Metagene profile of average H3K27ac, H2A.Z, and Brd4 enrichment at enhancer (Enh) regions. Enhancers were identified as regions enriched for H3K27ac that did not overlap with an annotated TSS. (**d**) Genome browser image depicting enrichment of H3K4me3 in HEK293T samples of the indicated cell number at a representative genomic region. A published ChIP-seq sample from ENCODE is provided for comparison. (**f**) Metagene profile of average H3K4me3 enrichment at the TSS region of annotated genes from the hg19 genome. Samples were obtained using the indicated number of cells. A published ChIP-seq sample from ENCODE is provided for comparison.

To evaluate the sensitivity of ACT-seq, we replicated the H3K4me3 data set using samples comprising varying cell numbers. Visual inspection of these data sets revealed that peaks were reliably detected using as few as 1,000 cells (**Fig. 1d**). This also held true for the pattern of average H3K4me3 enrichment at the TSS of genes (**Fig. 1e**). Consistent with these results, we observed strong correlation between statistically significant regions of enrichment in the H3K4me3 ChIP-seq data set and in each of the ACT-seq data sets obtained using different numbers of cells (**Supplementary Fig. 3**). These data support the reproducibility of ACT-seq data obtained from bulk samples of as few as 1,000 cells.

To map epigenetic marks in single cells, we adapted the ACT-seq method described above using a barcode multiplexing strategy. Permeabilized cells were divided volumetrically into 96 wells at a density of 5,000 cells per well. Each well was treated with a separate PA-Tnp complex carrying a unique combination of 5’ and 3’ sequence barcodes. After washing away any unbound complex, the cells were pooled and distributed into a second 96-well plate at a density of 18 cells per well using FACS sorting. The transposition reaction was initiated by addition of MgCl_2_ and terminated by addition of EDTA and proteinase. Library construction and amplification were performed separately in each well using a second set of distinct index barcodes that were unique to each well. The samples were then pooled, purified, and sequenced.

Using this indexing ACT-seq (iACT-seq) strategy, we mapped H3K4me3 enrichment in 1,246 individual cells (**Fig. 2**) and obtained about 2,500 unique reads per cell. Visual inspection of the mapped single-cell reads revealed that the read density predominantly clustered in the peaks of enrichment present in the bulk-cell ACT-seq and ChIP-seq data sets (**Fig. 2a**). Further, the pattern of average enrichment at the TSS regions of annotated genes was highly reproducible between the bulk-cell ACT-seq and single-cell ACT-seq data (**Fig. 2b**). Precision and sensitivity of the iACT-seq method compared favorably to Drop-ChIP^13^ (**Fig. 2c,d**), and visual evaluation of the mapped reads reveals that the precision of iACT-seq also compared favorably to ChIL-seq^14^ (**Supplementary Figure 4**). We found that peaks of statistically significant H3K4me3 enrichment were highly correlated between the iACT-seq data and the ENCODE ChIP-seq data set (**Fig. 2e,f**), in a similar manner to our analysis of the bulk-cell ACT-seq (**Supplementary Figure 3**). To determine whether our multiplexing strategy was successful at isolating individual cells, we repeated the iACT-seq experiment and substituted mouse pre-adipocyte cells in place of the HEK293T cells in 4 out of the 96 wells. Reads from this experiment were mapped to both the human hg18 and the mouse mm9 reference genomes (**Fig. 2g**). For all cells, we observed strong mapping to either the human or mouse genome, indicating that we successfully isolated single cells for analysis. We did not observe any samples with substantial mapping to both genomes, which would have been evidence that some samples erroneously comprised a combination of multiple individual cells. Based on our analyses, we conclude that iACT-seq is capable of efficiently mapping epigenetic marks in thousands of individual cells simultaneously, which takes one day of bench work.

**Figure 2:**
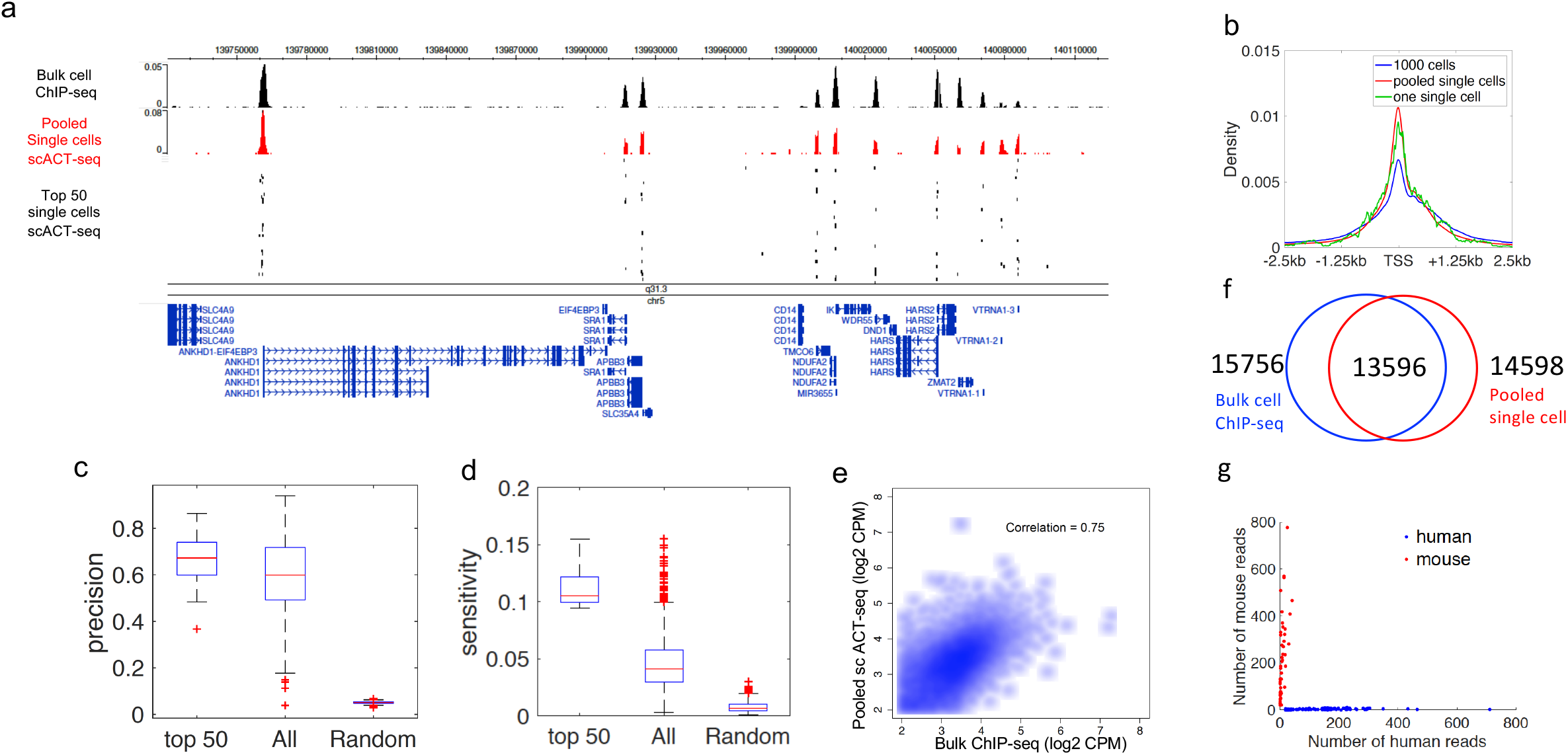
ACT-seq reproducibly maps epigenetic marks in single cells. (**a**) Genome browser image of H3K4me3 peaks from bulk-cell ChIP-seq (blue) and pooled scACT-seq (red). The mapped reads from the top 50 individual cells, as determined by read density, are plotted below the aggregate peaks. Each row represents an individual cell. (**b**) Metagene profile of average H3K4me3 enrichment at the TSS region of genes from the hg19 genome. Included samples are bulk-cell ACT-seq performed on 1,000 cells (blue), pooled scACT-seq (red), and one individual cell from scACT-seq (green). (**c-d**) Precision and sensitivity plots for the H3K4me3 scACT-seq data set. These values were calculated in the same manner as was done previously.^13^ (**e**) Scatter plots depicting the correlation in H3K4me3 peak enrichment in counts per million (CPM) between ENCODE bulk-cell ChIP-seq data (*x*-axis) and pooled scACT-seq data (*y*-axes). Peaks identified as enriched using both the ChIP-seq and scACT-seq methods were included. (**f**) Venn diagram indicating the numbers of significantly enriched H3K4me3 peaks with at least 1 bp overlap between a bulk-cell ENCODE ChIP-seq data set (blue) and pooled scACT-seq data (red). (**g**) Scatter plot depicting the number of reads that mapped to the human hg18 genome (*x*-axis) and mouse mm9 genome (*y*-axis) for a mixture of HEK293T cells (blue) and mouse pre-adipocytes (red) processed using iACT-seq.

In summary, we found that ACT-seq was capable of robustly mapping epigenomic features in bulk-cell samples including histone tail modifications, histone variants, and chromatin-binding proteins. Our data indicate that it is applicable to mapping epigenetic marks in a small number of cells and in single-cells. It takes only 5 to 6 hours to prepare libraries for a dozen epigenetic features and only one day of bench work to construct libraries for thousands of single-cells. Based on these attributes and its ease of use, ACT-seq presents an attractive alternative to existing techniques for profiling epigenetic marks in bulk and single-cell samples.

## METHODS

### Preparation of the PA-Tnp Transposon Complex for Analysis of Bulk Cell Samples

The expression vector containing the PA-Tnp fusion construct is available from Addgene under accession number 121137. Recombinant PA-Tnp protein was expressed in BL21-Gold(DE3) competent cells (Agilent cat # 230132) and purified using nickel beads. Quality and purity of the isolated protein was assayed using SDS-PAGE. 10 μM of recombinant PA-Tnp protein was incubated for 10 min at 25 °C in complex formation buffer (50 mM Tris pH 8.0, 150 mM NaCl, 0.05% Triton X-100, 12.5% glycerol) containing 50 μM of 5’ complex barcode and 50 μM of 3’ complex barcode. 1 μL of the appropriate antibody was added to 12 μL of prepared complex and incubated at 25 °C for 60 min. The antibodies used in this study were: anti-H3K4me3 (Millipore cat # 17-614), anti-H3K4me2 (Abcam cat # ab32356), anti-H3K4me1 (Abcam ab8895), anti-H3K27ac (Abcam ab4729), anti-H2A.Z (Abcam ab4174), anti-BRD4 (Bethyl cat # A301-985A100), and normal IgGs (Millipore cat # cs200581).

### Bulk-cell Chromatin Binding and Tagmentation

HEK293T cells were permeabilized using a 10 min incubation on ice in 500 μL of complex formation buffer per ~1 million cells. After permeabilization, all subsequent centrifugations of cells were performed using the following procedure: 1 min spin at 250 *g, rotate tubes 180°, repeat spin, leave ~10 μL of solution in the tube when removing buffer. Pelleted cells were washed once with 500 μL of complex buffer and suspended in another 500 μL of buffer. Aliquots of 1,000 cell equivalents were transferred volumetrically to clean microcentrifuge tubes and adjusted to 50 μL total volume using complex buffer. 5 μL of antibody-bound PA-Tnp complex was added to each 50 μL cell aliquot and incubated at room temperature for 60 min to allow chromatin binding. Unbound complex was removed using three washes performed as follows: pellet and suspend cells in 500 μL of wash buffer (50 mM Tris pH 8.0, 150 mM NaCl, 0.1% Triton X-100), rotate tube for 5 min at room temperature, repeat. Pelleted cells were rinsed with 500 μL of rinse buffer (50 mM Tris pH 8.0, 50 mM NaCl, 0.1% Triton X-100). The rinse buffer was removed and 90 μL of reaction buffer (50 mM Tris pH 8.0, 150 mM NaCl, 10 mM MgCl_2_, 0.1% Triton X-100) was added to the pelleted cells for a total volume of ~100 μL. The tubes were incubated for 30 min at 37 °C to allow transposition to occur. The reaction was terminated by addition of 4 μL of 0.5 M EDTA, 2 μL of 10% SDS, and 1 μL of 20 mg/mL Proteinase K followed by a 60 min incubation at 55 °C.

DNA was purified using phenol-chloroform extraction and ethanol precipitation. The DNA pellet was suspended in 10 μL of 10 mM Tris pH 8.0. PCR reactions for library preparation were performed by adding the following to the 10 μL DNA sample: 25 μL of Phusion High-fidelity PCR Master Mix (NEB catalog # M0531S), 1 μL of 5’ library index barcode, 1 μL of 3’ library index barcode, and 13 μL of nuclease-free water. Amplification was performed using an initial step of 72 °C for 5 min followed by 15 cycles of: 98 °C for 10 s, 65 °C for 30 s, 72 °C for 15 s, and a final extension step of 72 °C for 5 min. The PCR products were analyzed using agarose gel electrophoresis. Fragments of the desired size were excised and purified using a QIAquick Gel Extraction kit (Qiagen cat # 28506).

### Preparation of the PA-Tnp Transposon Complex for Analysis of Single Cells

Preparation of the PA-Tnp complex was performed in PCR tube strips totaling 96 wells, with each well receiving a unique combination of 5’ and 3’ complex barcodes. The following were mixed in each well: 1 μL of 1 μg/μL recombinant PA-Tnp protein, 0.75 μL of 10 mM 5’ complex barcode, 0.75 μL of 10 mM 3’ complex barcode, and 2.5 μL of 2X complex buffer. Tube strips were incubated at 25 °C for 10 min to allow complex formation. 1 μL of each of the 96 prepared PA-Tnp complexes were transferred to 96 fresh PCR tubes, carefully mixed with 0.8 μL of the desired antibody, and incubated at room temperature for 60 min.

### Single-cell Chromatin Binding and Tagmentation

1 million HEK293T cells and 0.1 million mouse pre-adipocytes were washed and permeabilized as described above. Aliquots of 10,000 permeabilized HEK293T cells were volumetrically dispensed into the wells. For the cell identification test experiment, four of the wells instead received 10,000 permeabilized mouse pre-adipocytes. The samples were mixed and incubated at room temperature for 60 min to allow chromatin binding. Unbound complex was removed by centrifuging the PCR tube strips for 3 min at 250 *g and carefully discarding all but ~10 μL of the solutions. 50 μL of complex buffer was added to each well, and the centrifugation and volume removal steps were repeated. The cell solutions were suspended and pooled together in a single microcentrifuge tube for a total of ~1 mL of combined volume. The tube containing the pooled cells was centrifuged for 1 min at 250 *g, rotated 180°, centrifuged again, and all but ~20 μL of solution was carefully removed. The cells were suspended in 200 μL of complex buffer and filtered using a cell strainer test tube (Corning cat # 352235). A FACS instrument was used to distribute the (un-stained) cells into 96 clean wells containing 18 cells each. The transposition reaction was initiated by adding 10 mM MgCl2 to each well in 10 μL final volume followed by a 37 °C incubation for 60 min. The reactions were terminated by addition of 12 mM EDTA, 0.1% SDS, and 1 μL of 1 AU/mL protease (Qiagen cat # 19157) to each well followed by a 60 min incubation at 55 °C. Each sample was transferred to a separate microcentrifuge tube and subjected to phenol-chloroform extraction followed by ethanol precipitation to purify the DNA. The samples were suspended in 10 μL each of 10 mM Tris pH 8.0 and transferred back to PCR tube strips.

For library preparation, 0.5 μL of 10 mM 3’ library index barcode was added to each of the 96 wells, with each well receiving a 3’ barcode with a unique sequence. Each well then received 10 μL of Phusion High-fidelity PCR Master Mix and 0.5 μL of 10 mM universal 5’ library index barcode. 18 cycles of PCR amplification were performed using the program described above. 10 μL of each PCR product were pooled and purified using three columns of a MinElute PCR Purification Kit (Qiagen cat # 28004) for a total elution volume of 30 μL. Fragments of the desired size were gel-purified as described above.

### Sequencing and Data Analysis

Paired-end sequencing was performed using an Illumina HiSeq 2500 platform. The resulting reads were mapped to the hg18 reference genome using Bowtie2.^15^ Data analysis and visualization was performed using custom R and Matlab scripts.

The TSS density profiles were computed using HOMER.^16^ H3K4me3 peaks were identified using SICER^17^ with following parameters: gap size = 200 bp and window size = 200 bp. The function “findOverlaps” from the R package GenomicRanges^18^ was used to compute the overlap between two sets of peaks. Spearman correlation was used to compare the read density between libraries. The ENCODE ChIP-seq data sets used for comparison were downloaded from GEO under accession numbers GSM2711409 (H3K27ac), GSM2711410 (H3K4me1), GSM2711411 (H3K4me2), and GSM945288 (H3K4me3)).

To determine whether individual cells were efficiently identified (**Fig. 2g**), we filtered out the possible doublets (samples potentially comprising more than one cell) and low-quality cells (with fewer than 500 reads) using the strategy described previously.^19^ In total, 1,246 single cells were obtained in this experiment. The mapping statistics of the single cells are included in **Supplementary Table 1**.

### Oligonucleotide Sequences

The names and sequences of oligonucleotides used in this study are available in Excel format in **Supplementary Table 2**.

## DATA AVAILABILITY

Next-generation sequencing data have been submitted to the Gene Expression Omnibus under accession number GSE125971 and will be made publicly available upon manuscript acceptance.

## ACKNOWLEDGEMENTS

We thank Dr. Kairong Cui for providing the NIH3T3 cells; the National Heart, Lung, and Blood Institute DNA Sequencing Core Facility for sequencing the libraries; and the National Heart, Lung, and Blood Institute Flow Cytometry Core facility for sorting the cells. The work was supported by Division of Intramural Research, National Heart, Lung and Blood Institute.

## AUTHOR CONTRIBUTIONS

BC, KZ, and WLK participated in writing the manuscript. KZ, BC, and QT designed the experiments and performed the bench work. WLK and BC performed the bioinformatics analyses and figure generation.

---

**Supplementary Figure 1:**
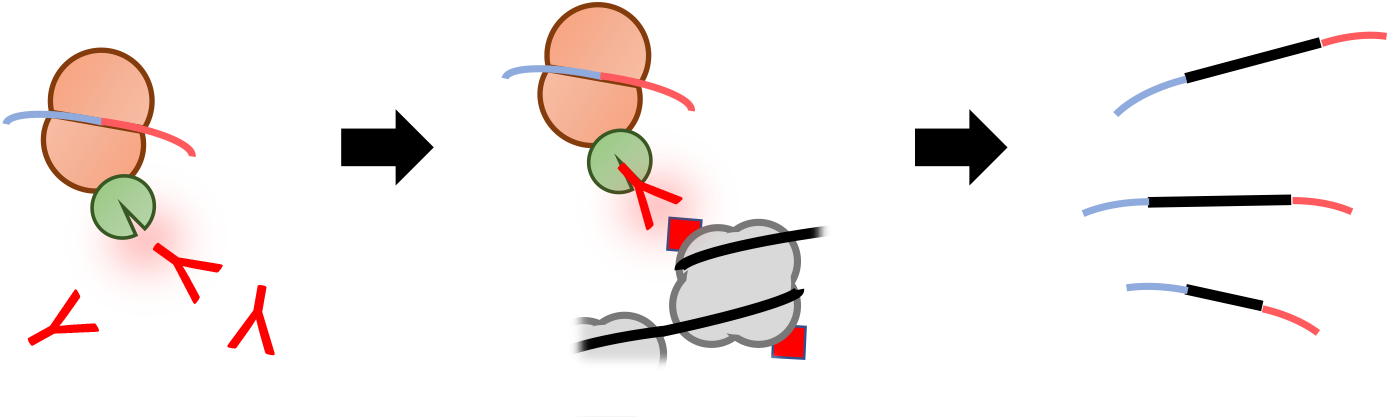
Diagram of the antibody-guided chromatin tagmentation strategy. Depiction of ACT-seq method steps from left to right: recombinant PA-Tnp transposome complex is bound to the selected antibody via the Protein A subunit (green); antibody-directed association of the transposome with the relevant antigen in chromatin; transposition and generation of tagged DNA fragments for sequencing.

**Supplementary Figure 2:**
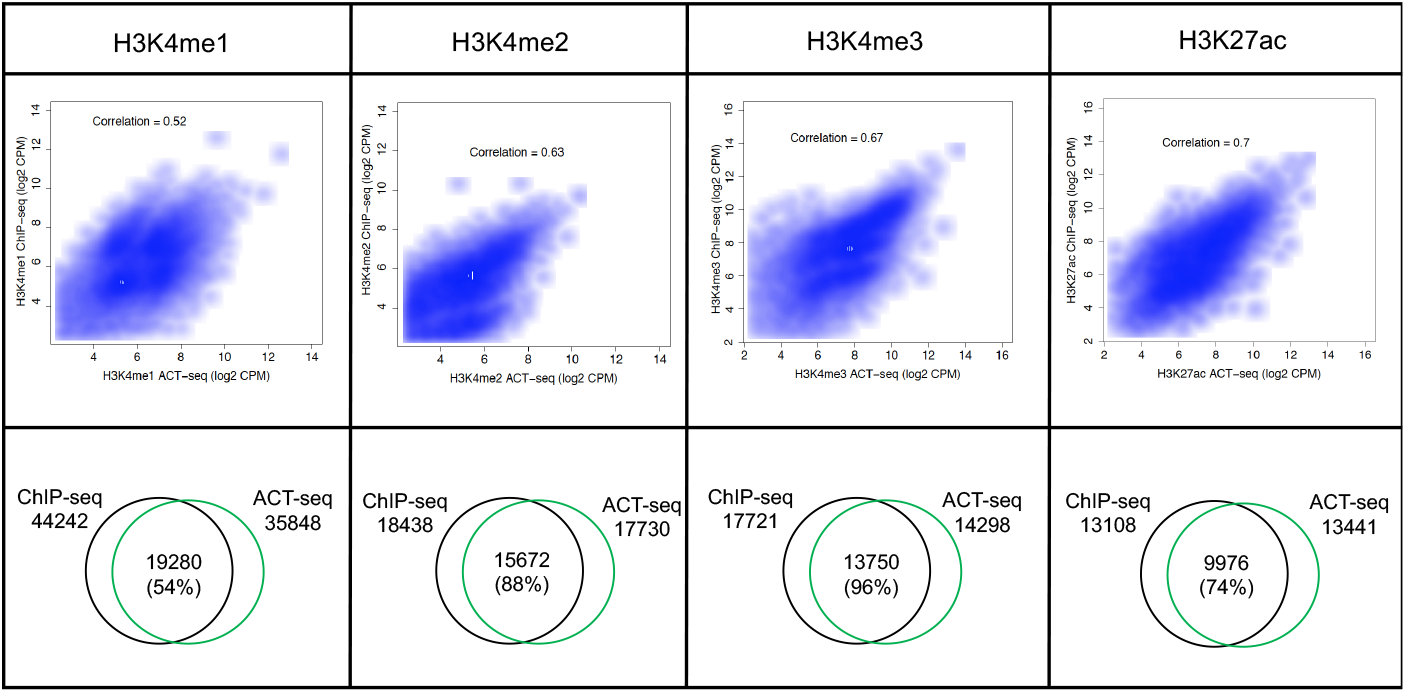
Enrichment detected using ACT-seq correlates with ChIP-seq data sets. (Top) Scatter plots depicting the correlation in peak enrichment in counts per million (CPM) between the ACT-seq data sets (*x*-axes) and ChIP-seq data sets (*y*-axes) for the indicated samples. Peaks identified as enriched using both the ChIP-seq and ACT-seq methods were included. (Bottom) Venn diagrams indicating the numbers of significantly enriched peaks with at least 1 bp overlap between the indicated ACT-seq samples (green) and published ENCODE ChIP-seq data sets (black). Percentages in parentheses represent the precision in measurements between ACT-seq and ChIP-seq (percentage = common peaks / total peaks in the given ACT-seq sample).

**Supplementary Figure 3:**
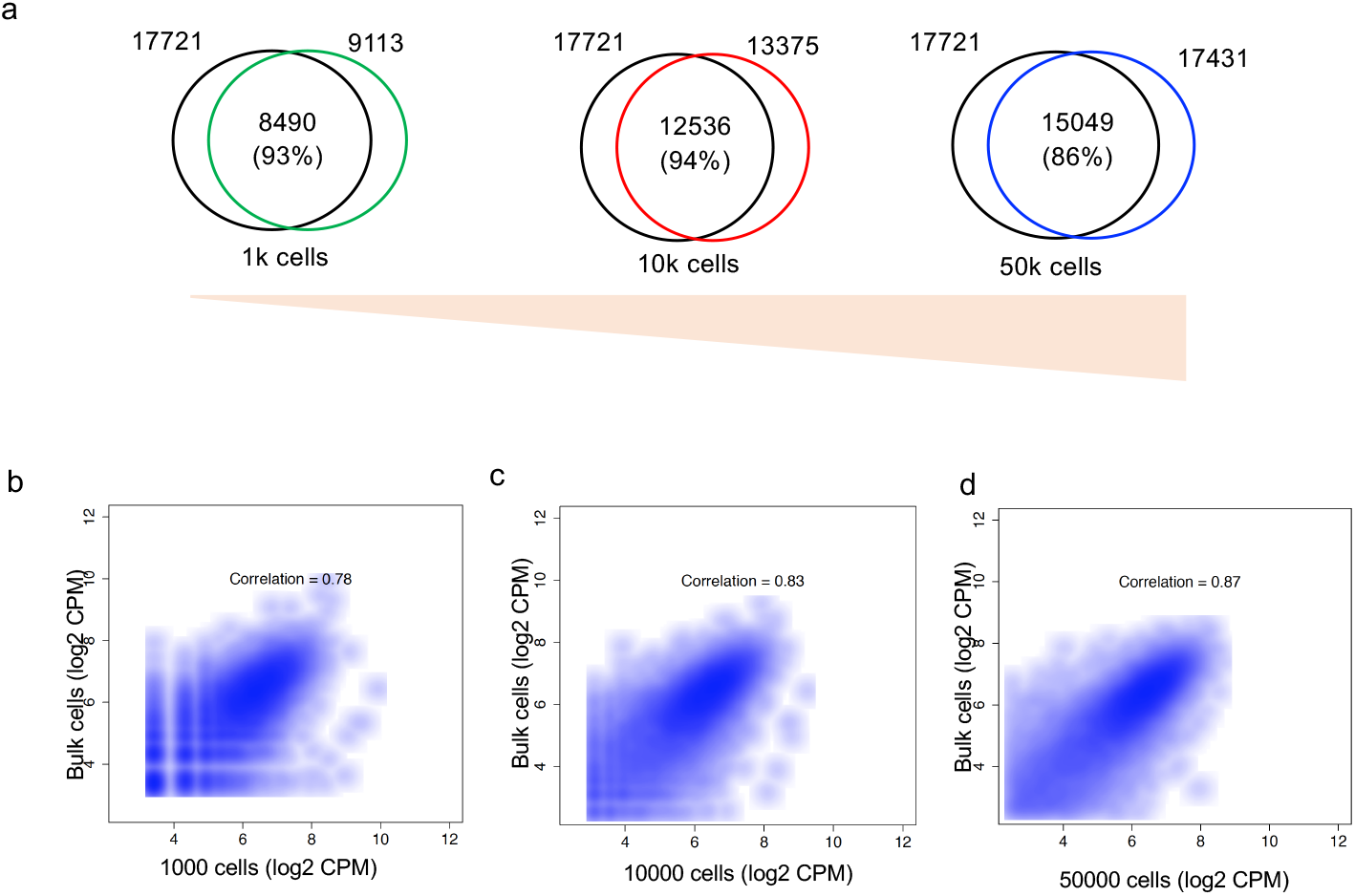
ACT-seq reproducibly detects H3K4me3 enrichment in as few as 1,000 cells. (**a**) Venn diagrams indicating the numbers of significantly enriched H3K4me3 peaks with at least 1 bp overlap between an ENCODE ChIP-seq data set (black) and ACT-seq samples obtained from the indicated numbers of cells. Percentages in parentheses represent the precision in measurements between ACT-seq and ChIP-seq (percentage = common peaks / total peaks in the given ACT-seq sample). (**b-d**) Scatter plots depicting the correlation in H3K4me3 peak enrichment in counts per million (CPM) between ACT-seq data sets obtained using the indicated numbers of cells (*x*-axes) and an ENCODE ChIP-seq data set (*y*-axes). Peaks identified as enriched using both the ChIP-seq and ACT-seq methods were included.

**Supplementary Figure 4:**
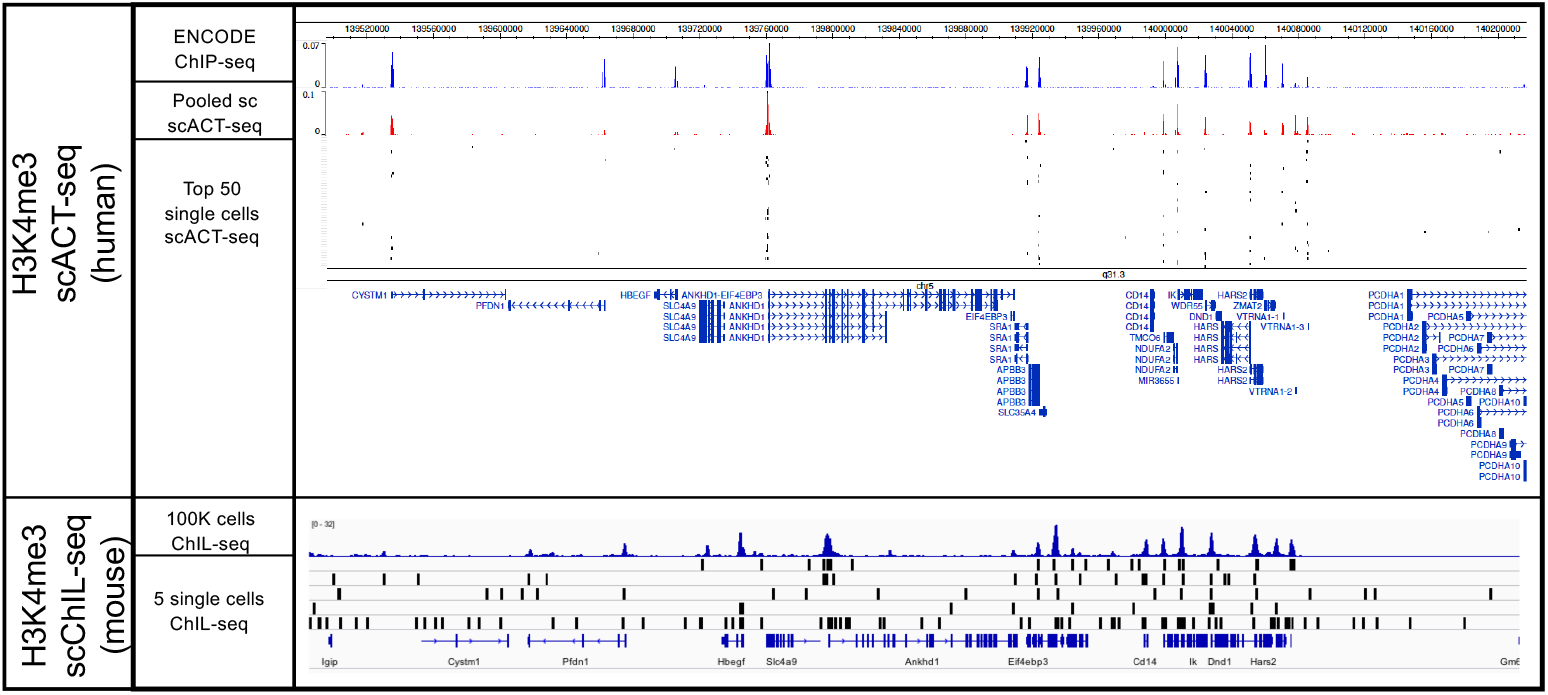
scACT-seq precision compares favorably with ChIL-seq. Genome browser image of scACT-seq (top) and ChIL-seq (bottom) data sets at representative genomic regions.

